# Unsupervised learning of haptic material properties

**DOI:** 10.1101/2021.02.25.432896

**Authors:** Anna Metzger, Matteo Toscani

**Author notes:** shared first authorship.

## Abstract

When touching the surface of an object, its spatial structure translates into a vibration on the skin. The perceptual system evolved to translate this pattern into a representation that allows to distinguish between different materials. Here we show that perceptual haptic representation of materials emerges from efficient encoding of vibratory patterns elicited by the interaction with materials. We trained a deep neural network with unsupervised learning (Autoencoder) to reconstruct vibratory patterns elicited by human haptic exploration of different materials. The learned compressed representation (i.e. latent space) allows for classification of material categories (i.e. plastic, stone, wood, fabric, leather/wool, paper, and metal). More importantly, distances between these categories in the latent space resemble perceptual distances, suggesting a similar coding. We could further show, that the temporal tuning of the emergent latent dimensions is similar to properties of human tactile receptors.

## Introduction

With our sense of touch we are able to discriminate a vast number of materials. We usually slide the hand over the material’s surface to perceive its texture (Lederman & Klatzky, 1987). Motion between the hand and the material’s surface elicits vibrations on the skin, which is the sensorial input mediating texture perception (Weber et al., 2013).

It was proposed that perceptual representations emerge from learning to efficiently encode sensorial input (Barlow, 1961; Olshausen & Field, 1996; see for a review: Simoncelli & Olshausen, 2001). For example, in color vision, the excitations of long- and middle-wavelength-sensitive cones are highly correlated. At the second stage of processing, still in the retina, a transformation into two color-opponent and a luminance channel achieves an efficient and decorrelated representation, akin to a principal components analysis of the input signals (Buchsbaum & Gottschalk, 1983; Zaidi, 1997; Gegenfurtner, 2003). Receptive field properties in the early visual pathway (Atick & Redlich, 1992; Olshausen & Field, 1996;1997) as well as the tuning properties of auditory nerve fibers (Lewicki, 2002; Smith & Lewicki, 2006) can emerge by efficiently encoding natural images or sounds respectively. Recently, it was shown that efficient coding could also explain the simultaneous development of vergence and accommodation as a result of maximizing coding efficiency of the retinal signals (Eckmann, Klimmasch, Shi, & Triesch, 2020). There is currently a lot of interest whether higher level representations can also be learned by efficient encoding of the retinal images (Fleming & Storrs, 2019).

Here we explore whether the perceptual representations of our haptic world can be learned by encoding vibratory signals elicited by the interaction with textures. We used a Deep Convolutional Autoencoder, a framework for unsupervised learning, to reconstruct the recorded signals from the LMT (Lehrstuhl für Medientechnik - Chair of Media Technology) haptic texture database (Strese, Boeck, & Steinbach, 2017, Figure 1A-B). This database contains acceleration recordings from free explorations of 108 everyday materials (Figure 1A-B).

**Figure 1.**
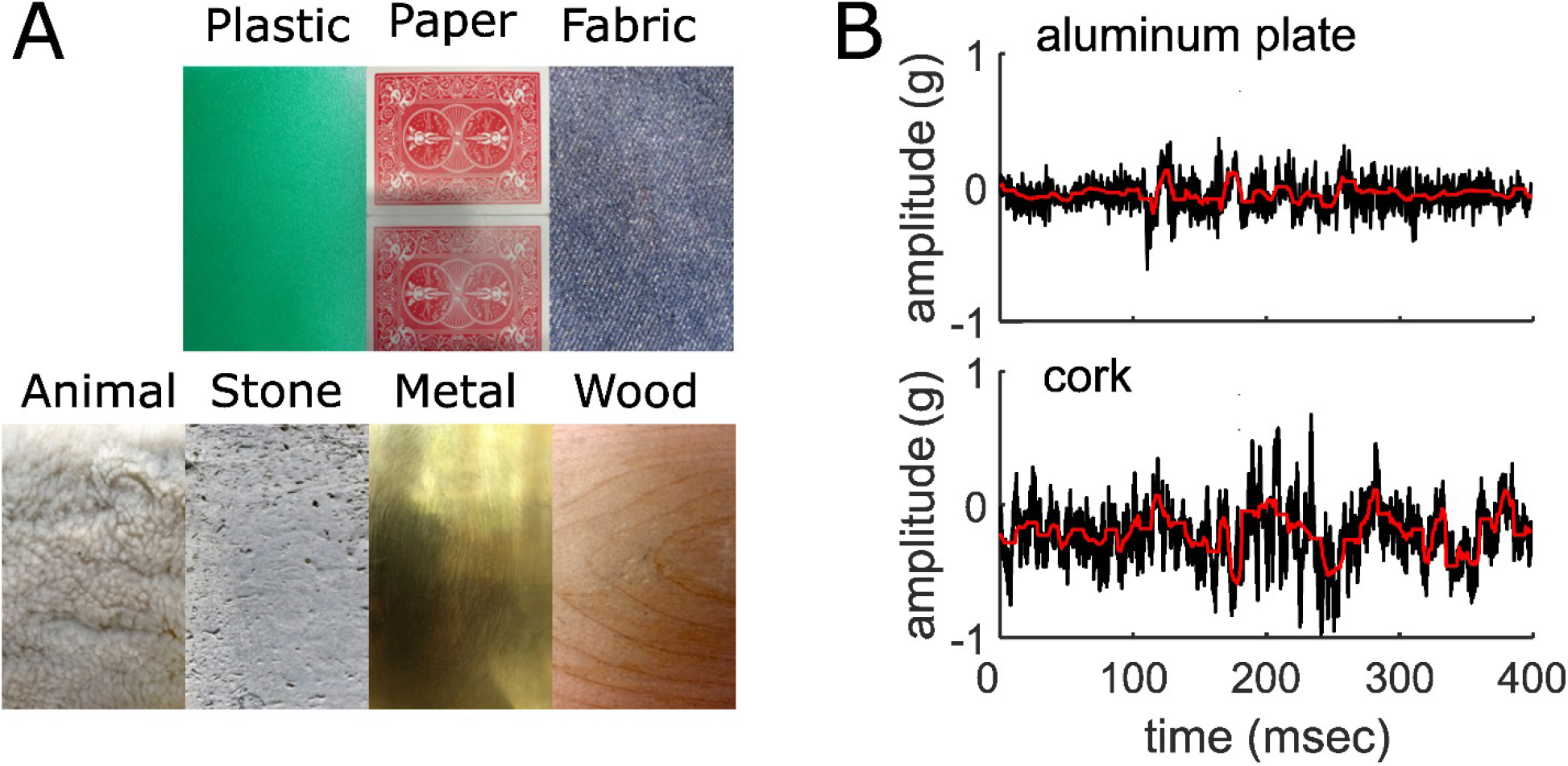
Materials and vibratory signals. A) Examples of materials from the LTM database. One example per category is shown. B) Original and reconstructed acceleration traces for two example materials. The black lines represent the original amplitude (y-axis) signals over time (x-axis), the red line the corresponding reconstructed signals.

We trained a Deep Convolutional Autoencoder to reconstruct the vibratory signals after compressing them into a 16 features latent representation. Then we related the latent representation to haptic material perception: we show that, based on the emerging latent representation, it is possible to classify the vibratory signals into the different material categories assigned by human participants. We computed the centroids of each category within the latent space, and showed that the distances between these categories resemble perceptual distances, measured with a rating experiment. These results suggest that the latent representation produced by unsupervised learning is similar to the information coding of the haptic perceptual system.

To interpret the dimensions of the latent space, we mimicked a physiology experiment. We generated a large number of sinusoids systematically varying in frequency, and computed the corresponding representation within the latent space. We observed that the temporal tuning of the latent dimensions is similar to properties of human tactile receptors responsible for perception of haptic textures (i.e. Pacinian (PC) and rapidly adapting (RA) afferents).

As a control, we repeated all analyses after training three additional network architectures. Since our results are replicated with four different networks, the similarities between the DNN latent representations and the perceptual representation are likely to be caused by efficient encoding rather than by the choice of the network’s architecture.

## Results

The Autoencoder learned a compressed latent representation to reconstruct the signals provided as input (Figure 1B). We evaluated reconstruction performance by correlating the input with the corresponding output signals and computing R^2^. Performance is computed for the train set (95% of the data), for the test set (remaining 5%) and for the full dataset. The reconstructed signals could explain 51% of the variance of the original signals (50.9% for the train set, 52.2 for the test set, and 51.2 for the full dataset). The learned latent representation is highly redundant: 99% of the variance of the latent representation of the full dataset can be explained with 2 Principal Components.

In order to relate this 2D representation (*latent PCs space*) to human perception, we labelled the materials of the LMT database according to a set of seven categories (Figure 1A) previously used to collect human judgments on material properties (Baumgartner, Wiebel, & Gegenfurtner, 2013). Linear classification could identify the correct category better than chance (i.e. 29%; empirical chance level given by a bootstrap analysis = 14%, with [9% 20%] 95% confidence interval). Classification based on the raw vibratory signals could be achieved only by extracting known features related to the psychophysical dimensions of tactile perception such as hardness or friction (Strese et al., 2017).

Some categories were easy to discriminate (Figure 2A), some more difficult. This is visible from the category averages in the *latent PCs space* (Figure 2B). For instance, *metal* is far apart from *animal* but close to *wood*.

**Figure 2.**
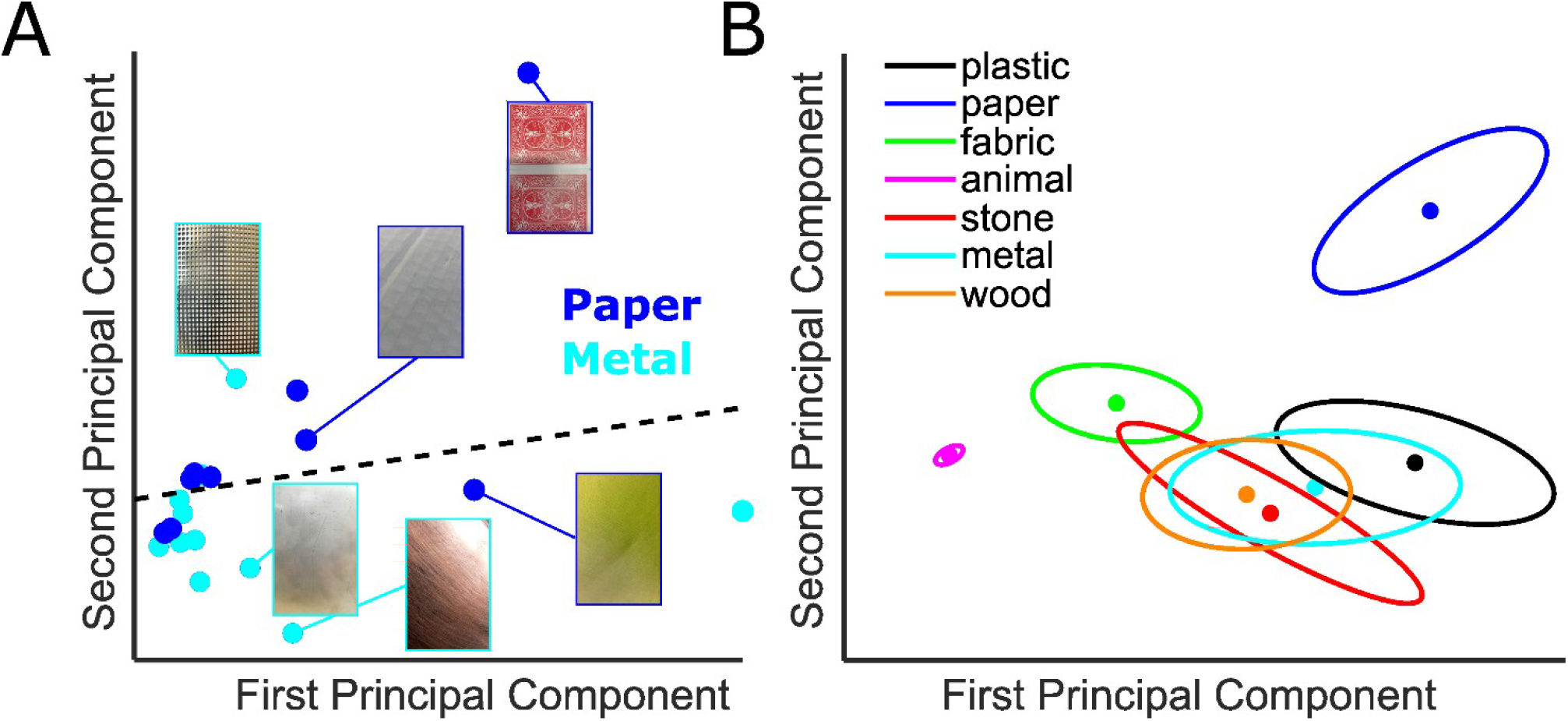
Latent space representation. A) Example of material category classification in the 2D latent PCs space. The blue data-points are the 2D representation of metals; paper textures in cyan. The icons represent examples of textures. The dashed line is the classification line learned by a linear classifier, which could tell apart the two categories with 75% accuracy. B) Centers of categories in the 2D latent PCs space. Each color represents a different category, as indicated in the legend. Dots represent the category average across materials, with the 2 dimensional errors represented by error ellipses (axes length given by 1/5 of the eigenvalues of the covariance matrix; axes direction corresponding to the eigenvectors).

Crucially, the distances between centers of categories in the *latent PCs space* correlate with perceptual similarity (Figure 3A – black data-points; N=21, r= 0.55, p=0.01), as inferred from Baumgartner et al.’s study (2013), indicating that the tactile representation of material properties within the compressed space learned by the Autoencoder resembles the perceptual representation.

**Figure 3.**
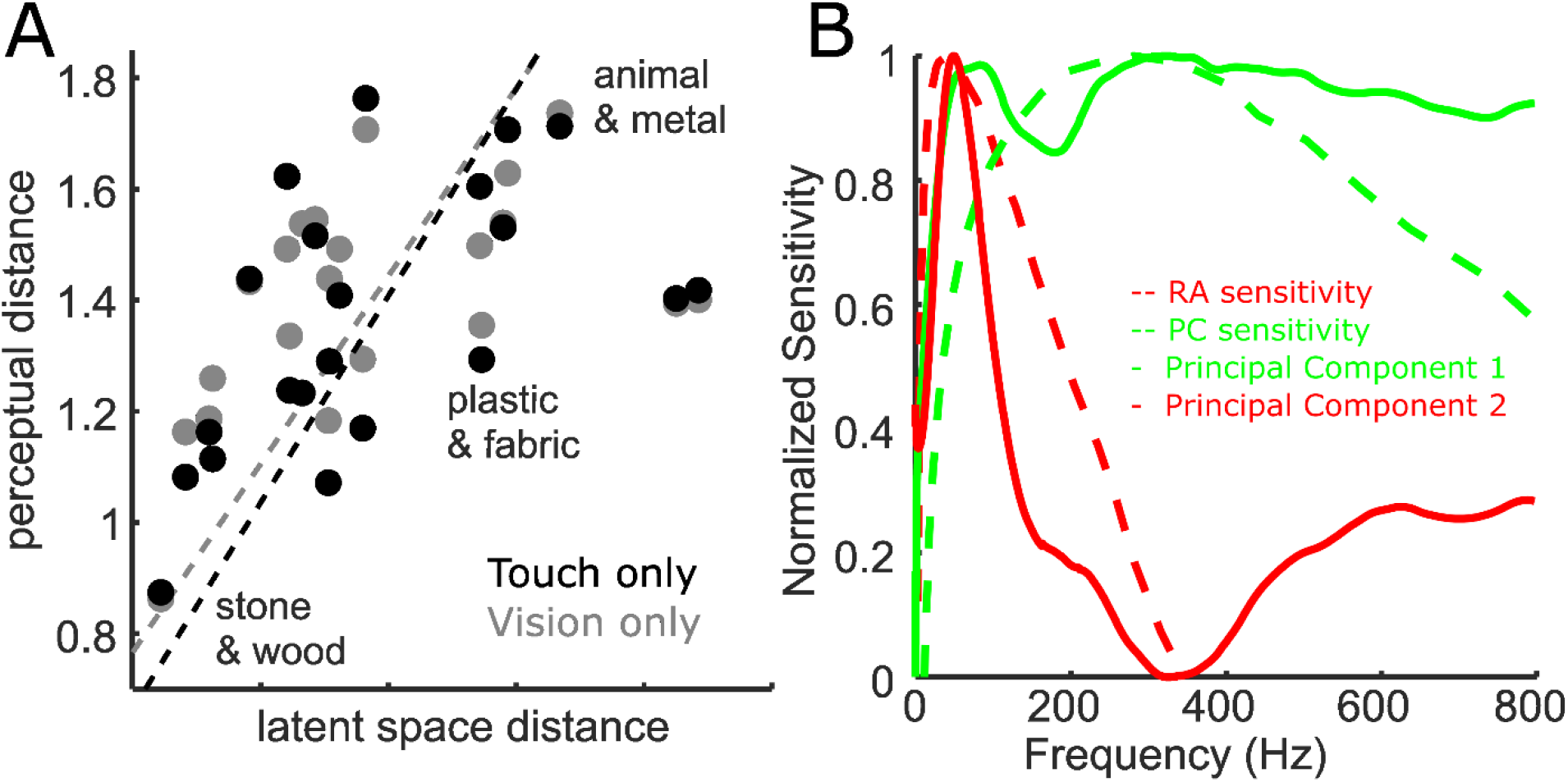
Relationship with perception. A) Relationship between perceptual distances (y-axis) and distances between categories in the 2D latent PCs space (x-axis). Distance with blind-folded observers is shown by black data-points; with vision-only by gray data-points. The black dashed line indicates the main principal component of the latent space distances and the perceptual distances for the blind-folded experiment, the gray line for the vision-only experiment. B) Comparison between temporal tuning of the two dimensions defining the latent PCs space (i.e. Principal Component 1 & 2 – continuous green and red lines, respectively), and the temporal tuning of the Pacinian (PC) and Rapidly Adapting (RA) afferents (green and red dashed lines, respectively). Tuning is given by sensitivity (y-axis) as function of temporal frequency (x-axis). RA is optimally tuned at 40-60 Hz, and the PC is optimally tuned at 250-300 Hz.

Baumgartner et al. (Baumgartner et al., 2013) repeated their rating experiment with only visual stimulation and again perceptual distances correlate with the distances between centers of categories in the *latent PCs space* (Figure 3A – gray data-points; N=21, r= 0.5, p=0.02). This is intriguing because visual and tactile signals originating from the same material are fundamentally different. However similar physical properties are likely to create similar patterns of covariations in the sensorial inputs. The similarity between the visual and the tactile representation of materials corroborates the idea that by learning to efficiently represent the sensory input, perceptual systems discover the latent variables responsible for the sensorial input (Fleming & Storrs, 2019), i.e, the systematic differences between materials.

Haptic textures are mainly sensed by the Pacinian (PC) and rapidly adapting (RA) afferents, which are highly sensitive to skin vibrations (Weber et al., 2013) and have known temporal tuning properties (Kandel et al., 2000). We probed the model’s temporal tuning by mimicking a physiology experiment: we presented the network with a large number of sinusoids systematically varying in frequency within the perceptually relevant range (Weber et al., 2013). We defined the temporal tuning of the *latent PCs space* as the responses to sinusoidal signals of different frequencies along each of its dimensions (Figure 3B).

The tuning curves of the two PCs are remarkably similar to the ones of the PC and RA afferents, suggesting that our sensors for texture (and material) properties have evolved to efficiently encode the statistics of natural textures as they are sensed through vibrations.

### Replication of the main results with different networks

We repeated all analyses after training three additional network architectures. Two are based on the original Convolutional Autoencoder, with a different number of features in the latent space (i.e. 8 and 32, instead of 16). The third deep neural network is a Variational Autoencoder (VAE), with 16 latent dimensions.

All models could reconstruct the original signals, to some extent. Performance decreases with compression rate (Table 1).

**Table 1.** Reconstruction accuracy for the different networks. Accuracy is measured as the variance of the original signals explained by the reconstructed signals (R^2^). R^2^ is computed for each sample; medians across samples are reported in the table. Accuracy is computed for the train set (95% of the data), for the test set (remaining 5%) and for all data-set together. The last column indicates the network compression rate, i.e. the ratio between the dimensionality of the latent representation and the one of the original signals, expressed as percentage. Accuracy is lower with higher compression.

|  | ***R-squared*** |  |  | ***Compression rate*** |
| --- | --- | --- | --- | --- |
|  | ***train set*** | ***test set*** | ***full dataset*** |  |
| ***8 Features*** | 47.9% | 48.4% | 48% | 3% |
| ***16 Features*** | 50.9% | 52.2% | 51.2% | 6% |
| ***32 Features*** | 57.9% | 59.4% | 58.1% | 12% |
| ***VAE*** | 12% | 12.7% | 12.1% | 0.03% |

The *latent PCs space*, computed for all networks as for the original DNN, allows for classifying material categories: accuracy is 29% with the 32 Features network (empirical chance level given by a bootstrap analysis = 15%, with [9% 21%] 95% confidence interval), 23% with the 8 Features network (empirical chance level given by a bootstrap analysis = 15%, with [9% 21%] 95% confidence interval), and 21% with the VAE network (empirical chance level given by a bootstrap analysis = 14%, with [8% 21%] 95% confidence interval).

The perceptual distances inferred from the ratings from the blind folded observers correlate with the distances within the *latent PCs space* of all networks (Figure 4): Pearson’s r = 0.57, 0.55 and 0.45, for the 32 features, the 8 Features and the VAE networks, respectively (p-values= 0.008, 0.01, 0.04, respectively).

**Figure 4.**
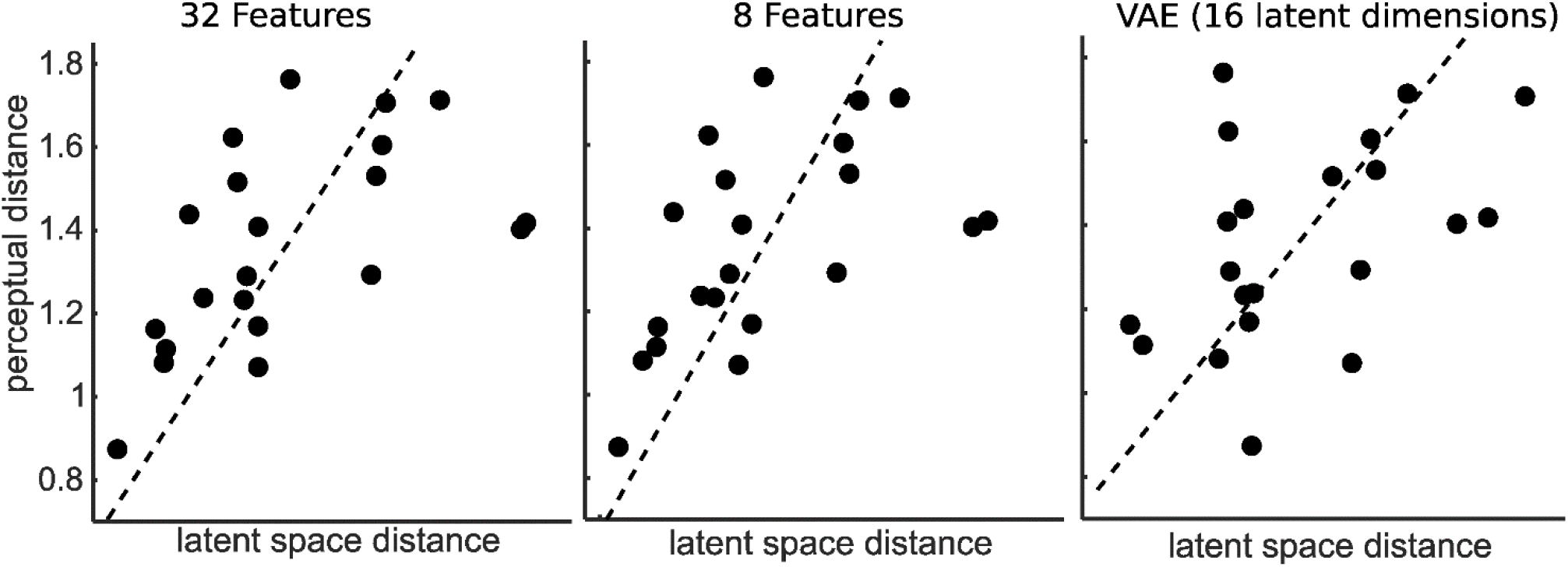
Relationship between perceptual distances (y-axis) and distances between categories in the latent PCs space (x-axis), for the control networks. The distance inferred from category ratings with blind-folded observers is shown by black data-points; with vision-only by gray data-points. The black dashed line indicates the main principal component of the latent space distances and the perceptual distances for the blind-folded experiment, the gray line for the vision-only experiment. Results are shown for the 32 Features network the 8 Features network and the VAE, from the left to the right panel, respectively.

Figure 5A shows the positions of the centers of categories in the space individuated by the two first principal components of the DNNs features space (or of the latent dimensions of the VAE). The VAE does not explicitly represent time in its latent dimensions. Therefore, PCA was applied directly on the 16 latent dimensions and the two main components were considered for the analysis. We used the procrustes transformation to “superimpose” the centres of the categories for the 8, the 32 Features and the VAE networks to the ones of the original 16 Features network. Centers of categories within the latent space of the 8, 16, and 32 Features networks are nearly overlapping, indicating that these networks learned a similar representation despite the different compression rate.

**Figure 5.**
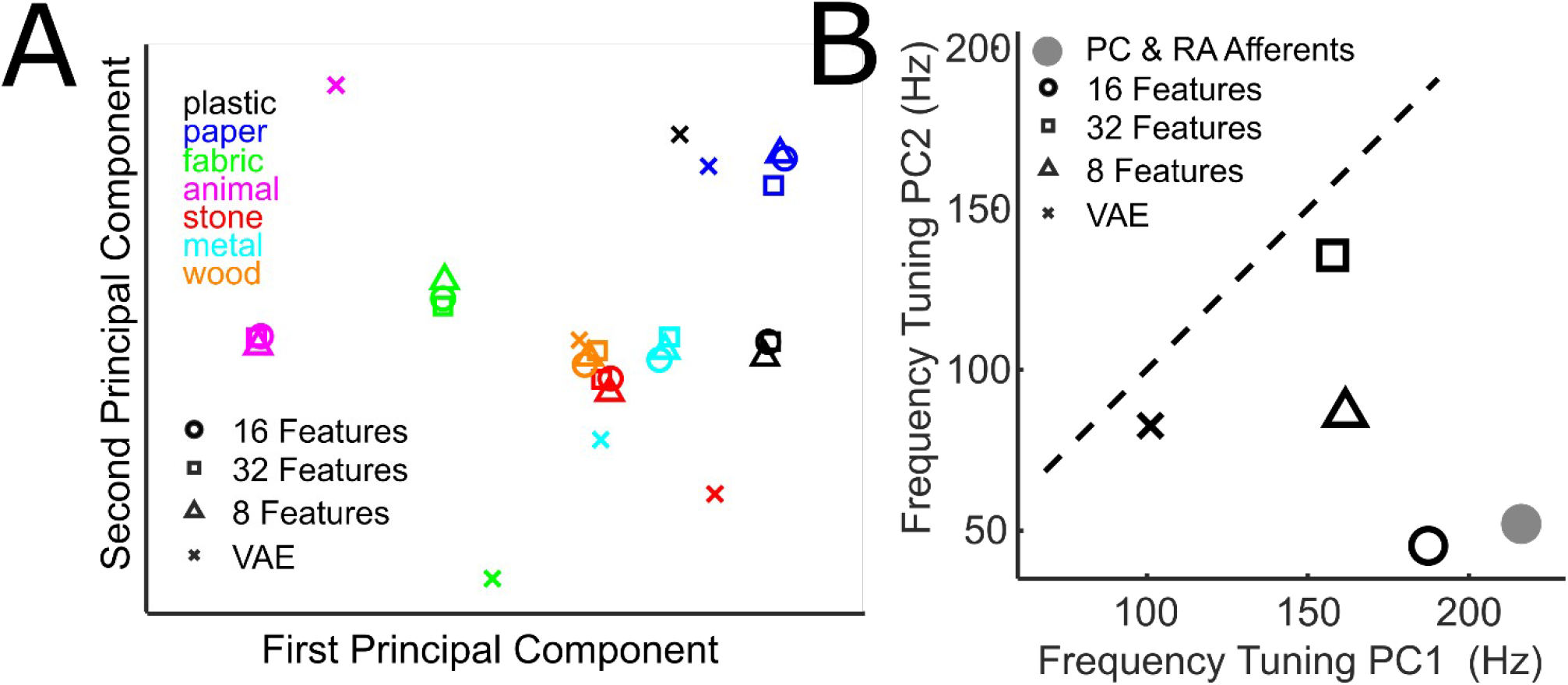
Comparison of the categories representation and temporal tuning of the latent PC space of all networks. A) Centers of categories in the latent PCs space. Each color represents a different category, as indicated in the legend. Different symbols represent the category average across materials for the different networks, i.e. squares for the 32 Features network, circles for the 16 Features network, triangles for the 8 Features network, squares for the 32 Features network and crosses for the Variational Autoencoder. B) Comparison between temporal tuning of the two dimensions defining the latent PCs space. Tuning of the first principal component (PC1) on the x-axis, of the second principal component on the y-axis. Different symbols indicate the different networks (see legend). The gray circle indicates the temporal tuning of the Pacinian (PC, x-axis) and the Rapidly Adapting (RA, y-axis) mechanoreceptors. The dashed line is the unity line.

The VAE shows clear differences with the other networks, presumably because of the much higher compression rate (the latent representation is 200 times smaller than the one of the 16 Features network), or the different architecture (see methods). However, the distances between centers are very similar to the ones computed within the latent spaces of the other networks (Pearson’s r= 0.861, 0.86, and 0.858; for the 8, 16 and the 32 Features network, respectively). This is consistent with the correlation we found between the distances within the VAE *latent PC space* and perceptual distances.

For all networks, one of the two dimensions of *latent PCs space* is tuned to relatively low temporal frequencies and the other to higher frequencies. A measure of spatial frequency tuning is computed by weighting each frequency by the corresponding sensitivity and averaging across frequencies. Tuning is computed for the first Principal Component and for the second Principal Component of the *latent PCs space*, for all networks, and for the two afferents. Although there are differences between the four networks, PC1 is always tuned to higher frequencies than PC2 (Figure 5B).

## Discussion

We used unsupervised learning to reconstruct the vibratory signals that constitute the very input of the haptic system, based on a highly compressed representation. Such representation shares similarities with the perceptual representation of natural materials: it allows for classification of material categories, as provided by human judgements, and crucially, distances between these categories resemble perceptual distances. Furthermore, the temporal tuning of the dimensions of the learned representation is similar to properties of human tactile receptors. These similarities suggest that the computations involved in touch perception evolved to efficiently encode the input, yielding perceptual representations that allow to distinguish between different materials. This may be because a way to efficiently represent the vibratory patterns is to discover latent variables that create systematic differences between them, i.e. the systematic physical differences between materials.

A similar idea has been proposed for visual perception of material properties (Fleming & Storrs, 2019). The challenge for vision is that the information in the retinal image (proximal stimulus) is insufficient to recover the properties of the world (distal stimulus) that caused it (Anderson, 2011). In fact, multiple causes are usually responsible for a proximal stimulus, e.g. illumination and reflectance spectra are confused within the reflected light. To solve this ambiguity, it is proposed that we learn to represent the dimensions of variation in the proximal stimuli, which arise from the systematic effects of distal stimuli (Fleming & Storrs, 2019), rather than learning to directly estimate the distal properties of the world, as predicted by the *inverse optic*s approach (Marr, 1982; Poggio & Koch, 1985; Pizlo, 2001). Our results support the hypothesis that efficiently encoding the proximal stimuli is the way sensory systems develop perceptual representations.

Similar to the ambiguities in the visual input, a challenge for touch perception is that the temporal frequency content of the input vibratory signals depends both on the surface properties and the exploration speed. Nevertheless, we can recognize different materials regardless of the speed of the hand movements used to explore them. This ability of the haptic system can be referred to as speed invariance (Boundy-Singer, Saal, & Bensmaia, 2017). Our classification analysis could discriminate the material categories even though exploration speed was not controlled (Strese, 2017). This indicates a certain degree of speed-invariance, consistent with the fact that human roughness judgments can be predicted from non-speed invariant responses of the PC and RA afferents (Weber et al., 2013), presumably because of limited variability in exploration speed. Speed-invariance can be implemented at a later stage by dividing normalizing the responses by the movement speed, yielding a representation in spatial coordinates. For this, the haptic system would need a precise and robust measure of the movement speed. An alternative mechanism for speed invariance could be similar to the one mediating (auditory) timbre constancy: while changes in exploration speed affect the frequency composition of the receptors’ spiking responses, the harmonic structure remains relatively constant (Manfredi et al., 2013).

However, it is possible that observers adjusted their exploration movements to the best speed for texture recognition, yielding a certain level of speed invariance. It is known that exploratory behavior is optimized to maximize information gain (Lederman & Klatzky, 1987; Najemnik & Geisler, 2005; Kaim & Drewing, 2011; Toscani, Valsecchi & Gegenfurtner, 2013). Specific to exploration speed, it is shown that when participants were asked to haptically discriminate spatial frequencies of gratings, low performing participants could improve their performance by adjusting the scanning velocity to the one used by better performing participants (Gamzu & Ahissar, 2001). In fact, the statistics of the sensory input are shaped by exploratory behavior, which therefore may contribute to efficient encoding, consistent with the recently proposed “active efficient coding” theory (Eckmann et al., 2020).

Here we show that efficient encoding of the raw vibratory signals (i.e. measured at the tool with which the surface texture is explored) produces a representation similar to the signals recorded from the afferents responsible for texture perception (i.e. PC and RA). This does not imply that the different auto encoders we tested only mimic the function of the Pacini and Meissner corpuscles innervated by the PC and RA afferents, because, before the receptors, the signal might already be compressed by the biomechanic properties of the hand (Shao, Hayward & Visell, 2020).

Control experiments suggest that our results are likely to be caused by efficient encoding rather than by the choice of the network’s architecture, as they are replicated with four different networks.

Because of constraints on perceptual systems, like the number of neurons and the metabolic cost of neural activity (Lennie, 2003), receptor input is likely to be compressed and encoded as efficiently as possible to maximize neural resources. Our results suggest that efficient coding forces the brain to discover perceptual representations that relate to the physical factors responsible for variations between signals.

## Methods

### Deep Neural Network

The network used for the analysis described in the manuscript is a relatively simple deep convolutional Autoencoder. The *encoder* encodes the 47872 time samples of each vibratory pattern into a latent representation consisting of 187 time samples per 16 features (i.e. *code*). This is done by means of four 1D convolutional layers, each of them with kernel size equal to three time samples.

The size of the representation in time is progressively reduced by means of a max pooling operation, so that the kernel size is relatively increased for deeper layers. Specifically, after the input layer, the first convolutional layer takes as input signals of 47872 time samples and outputs 47872 times samples for each of 64 features. The max pooling operation reduces the time samples to 11968. The second convolutional layer takes as input 11968 time samples per 64 features and outputs 11968 time samples per 32 features. Again, max pooling reduces the times samples; this time to 2992. The third convolutional layer takes as input 2992 time samples per 32 features and outputs 2992 time samples per 16 features. Max pooling reduced the time samples to 748. The fourth convolutional layer takes as input 748 times samples per 16 features, and outputs the same size representation.

Finally, max pooling reduces the time samples to 187. This output, consisting of 187 time samples for 16 features, is the most compressed representation within the network, i.e. the *code*, and constitutes the bottleneck of the Autoencoder.

The *code* is decoded by the decoder, to reconstruct the input signals. The first convolutional layer of the decoder takes as input, and outputs 187 time samples per 16 features. The Upsampling operation increases the number of time samples from 187 to 748. This is done by repeating each temporal step 4 times along the time axis. The second convolutional layer takes as input, and outputs 748 time samples for 16 features. Upsampling increases the time samples from 748 to 2992. The third convolutional layer takes as input 2992 time samples per 16 features, and outputs 2992 time samples per 32 features. Upsampling increases the time samples from 2992 to 11986. The fourth convolutional layer takes as input 11986 time samples for 32 features, and outputs 11986 time samples for 64 features. Upsampling increases the time samples from 11986 to the original size (i.e. 47872). The fifth and last convolutional layer takes as input 47872 time samples per 64 features and outputs 47872 time samples. Thus, input and output have the same size and can be directly compared within the loss function.

The activation function (**Π**) of all convolutional layers is the rectified linear unit (Equation 1), with the exception of the last layer of the decoder, for which it is a sigmoid function (Equation 2).

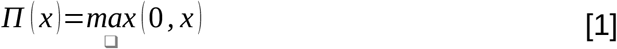

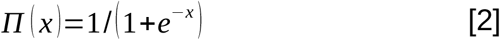

Training consisted of 50 epochs, to minimize the *mean absolute error (MAE)* loss function (equation 1).

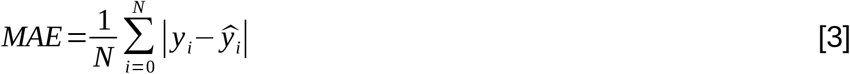

With *y*_*i*_ the *i* vibratory signal of the training set (consisting of N signals), and 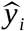 the corresponding reconstructed signal.

The network was adapted from (Chollet, 2016) to work with one-dimensional data rather than images. The implementation and training of the network was done through the Keras deep learning framework (Chollet, 2015) via its Python interface.

### Control Neural Networks

Two DNNs are based on the architecture of the network used for the analysis in the main text, with a different number of features in the latent space (i.e. 8 and 32, instead of 16). We increased the Features to 32 by changing the third and the fourth convolutional layers of the encoder, so that the former takes 2992 time samples per 32 features as input, and outputs 2992 time samples per 32 features instead of 16, and the latter takes as input, and outputs 748 time samples per 32 features.

Therefore, the *code* consists of 187 time samples per 32 features. The first convolutional layer of the decoder is modified so that it takes as input, and outputs 187 time samples per 32 features. The second, takes as input, and outputs 748 time samples per 32 features. The last difference with the original architecture is that the third convolutional layer of the decoder takes as input 32 features instead of 16.

We reduced the Features from 16 to 8 by changing the fourth convolutional layer of the encoder so that it outputs 8 features instead of 16, i.e. the *code* consists of 187 time samples for 8 features. Consistently, we changed the first layer of the decoder so that it takes as input and outputs 8 features, and the second layer takes 8 features as input and outputs 16 features. The differences in the number of features for the bottleneck layer, imply a different compression rate: 3% and 12%, for the 8, and 32 features networks, respectively.

The third deep neural network is a Variational Autoencoder (VAE), with 16 latent dimensions. With this architecture, the encoder represents each input vibratory pattern *i* with two parameters for each dimension of the latent space: the mean and the standard deviation (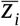 and *σ*_*i*_, respectively) of a Gaussian distribution that is assumed to generate the data. Then, one z point is randomly sampled from the corresponding Gaussian distribution (Equation 4).

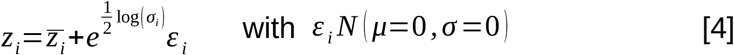

Thus, each vibratory pattern is associated to a point in the 16 dimensional latent space, and time is no longer represented, yielding higher compression than with the other architectures (0.03% of the input data). The decoder maps these latent space points back to the original input data.

The parameters of the model are trained by minimizing a custom loss function *L* (Equation 5) consisting of a reconstruction term (*BCE* – binary cross-entropy; Equation 6), a measure of error between the *N* decoded samples (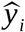 with *i ∈* [1 *N*]) and the initial inputs (*y*_*i*_ with *i ∈* [1 *N*]), and a regularization term (RT; Equation 7), that expresses divergence between the learned latent distribution and the assumed Gaussian distribution.

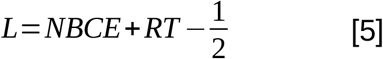

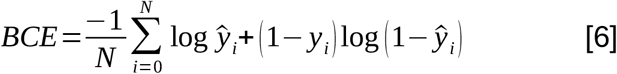

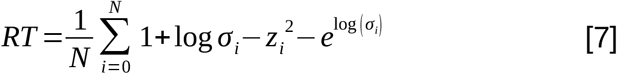

Again, the network was adapted from (Chollet, 2016 – Convolutional Variational Autoencoder) to work with one-dimensional data rather than images. For training the Variational Autoencoder we used data augmentation. For this we used random cropping: we sampled 20000 sub-portions of the original 4 seconds signals, by randomly selecting each time one sample from the original set and from this sample a random sub-portion of 150 milliseconds.

After the input layer, the first convolutional layer of the encoder takes as input 1500 time samples and outputs 750 time samples per 32 features. The time samples are reduced by half by setting the stride length of the convolution to 2 time samples. The second convolutional layer takes as input 750 time samples per 32 features and outputs 375 time samples per 64 features (again, the stride length is set to 2). After, a flatten layer converts the 375 time samples and 64 features into a 375*64 = 24000 samples representation. This representation is given as input to a fully connected (Dense) layer with 16 units. This is then given as input to two different fully connected layers, with 16 units each: the “z mean” layer and the “z variance” layer. Finally, a custom “sampling” layer samples one z point per unit, from a Gaussian distribution with mean and variance given by the activation of the 16 units in the “z mean” and the “z variance” layers, respectively. These 16 dimensional z points constitute the compressed representation (code).

The code is decoded as follows: z points are given as input to a fully connect layer that outputs a 24000 samples representation. This is reshaped into a 375 time samples per 64 features representation. The next three layers use transposed convolution to upsample the input feature maps to restore the size of the original signals. First the signal is converted from 375 time samples per 64 features, to 750 time samples per 64 features. Then, to 1500 per 32 features; finally to 1500 time samples.

Again, the activation function of all convolutional layers is the rectified linear unit (Equation 1), with the exception of the last layer of the decoder, for which it is a sigmoid function (Equation 2).

### Vibratory signals

We used acceleration recordings of human free hand explorations of 108 different materials with steel tool tip which are openly available in the LMT Database (Strese et al., 2017). Each vibratory signal consists of 5 seconds recordings. For each of the 108 material textures, 20 vibratory signals were acquired from 11 different people at 10000Hz temporal resolution, therefore each of them consisting of 50000 time samples. We cut the first 212.8 ms to remove the portion of the signals, which is affected by the onset of contact between the tool and the material surface. Frequencies below 10Hz were ascribed to exploratory hand movements and therefore were filtered out (Strese et al., 2017). Categories are taken from (Baumgartner et al.,2013) and assigned to the 108 textures of the database by two independent observers who agreed for all samples.

### Latent representation

The Autoencoder compresses the 47872 time samples of each vibratory pattern into 187 time samples represented per 16 features, i.e. down to 6% of the original. After reducing the 16 features to 2 Principal Components, for all analyses we represented each vibratory pattern as a point within the *latent PCs space* by averaging across time, since we were interested in the feature space. We also averaged across multiple measures of each material. This choice, is however not crucial, because classification of categories is also possible based on the individual signals (see *linear classification* section).

### Linear classification

Classification was done based on the *latent PC space* dimensions, by iteratively leaving out one material per category, training the classifier on the remaining signals and computing performance on the left out samples. Some categories include more materials than others, i.e. data is unbalanced for classification. We overcame this limitation by oversampling vibratory patterns to match their number across categories. As a conservative strategy for hypothesis testing we used bootstrap analysis to assess the significance of all classification results. For that, we repeated the classification analysis 5000 under the null hypothesis of no relationship between the points in the latent PC space and the material categories, i.e. we shuffled the category labels. This produced the accuracy distribution under the null hypothesis of chance-level classification; the 95% confidence interval was computed by reading out the 2.5^th^ and the 97.5^th^ percentiles of the distribution. The empirical chance level corresponded to the mean of that distribution.

We repeated the classification analysis based on the individual signals (rather than averaging across the different recordings for each material), by iteratively leaving out one vibratory signal per category, training the classifier on the remaining signals and computing performance on the left out signals. Accuracy (27%) is above chance (empirical chance level given by a bootstrap analysis = 14%, with [13% 16%] 95% confidence interval), indicating that samples of materials of different categories clustered together in the latent PCs space.

### Perceptual distances

Baumgartner et al. (2013) asked participants (blind-folded in the “touch only”, or with vision but no touch in the “vision only” condition) to rate ten properties of 84 different materials from seven material categories (i.e. “plastic”, “paper”, “fabric”, “animal” – i.e. leather or fur, “stone”, “metal”, “wood”), then computed correlations of ratings between categories. Perceptual distance (*δ*) is computed from the correlation coefficient (*ρ*) with the following equation:

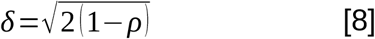

### Temporal tuning

We fed the network with 830 sinusoids with frequency ranging from 0 to 800 Hz, and computed their representation in the *latent PCs space*. Temporal tuning is given by the normalized (range is forced from 0 to 1) model responses, as a function of temporal frequency. For the tactile afferents, sensitivity is computed as normalized inverse of the discrimination thresholds from (Kandel et al., 2000).

## Acknowledgements

This work was supported by Deutsche Forschungsgemeinschaf (DFG, German Research Foundation) – project number 222641018 – SFB/TRR 135, A5 & A8. We are grateful to Karl Gegenfurtner and Knut Drewing for helpful discussions and comments.

## Data availability

The analysis code is publically available. The code for processing the vibratory signals from database (LMT) and the python code for training the DNNs and saving the reconstructed signals and the latent representations is available at github (https://github.com/matteo-toscani-24-01-1985/material_categories).

